# Displaced myonuclei are attributable to both resident myonuclear migration and stem cell fusion during mechanical loading in adult skeletal muscle

**DOI:** 10.1101/2025.10.10.681647

**Authors:** Nathan Serrano, Pieter Jan Koopmans, Kevin A. Murach

## Abstract

Non-peripheral myonuclei are characteristic of skeletal muscle pathology and severe injury but also appear after exercise and with aging. Displaced myonuclei are typically attributed to the activity of muscle stem cells, or satellite cells. We sought to address whether displaced myonuclei in adult skeletal muscle are exclusively from an exogenous source such as satellite cells or can result from resident myonuclear migration. To address this question, we used a murine recombination-independent muscle fibre-specific doxycycline-inducible fluorescent myonuclear labelling approach, EdU stem cell fate tracking, two durations of muscle mechanical overload (MOV, 3 days and 7 days), and fluorescent histology. Our findings show that: 1) displaced myonuclei emerge early during MOV, 2) resident myonuclear movement occurs rapidly during MOV, and 3) the contribution of resident versus exogenous displaced myonuclei depends on MOV duration, fibre type, and fibre size. These observations provide fundamental insights on myonuclear motility in response to stress *in vivo* and reframe our understanding of how a recognized feature of mammalian skeletal muscle can emerge in response to mechanical loading.

**Summary:** Recombination-independent muscle fibre-specific doxycycline-inducible fluorescent myonuclear labelling in adult mice unambiguously reveals how resident myonuclei relocate rapidly during stress and contribute to the appearance of displaced myonuclei.

## Introduction

In healthy adult skeletal muscle, nuclei in syncytial muscle fibres (myonuclei) are situated on the periphery of the cell. This peripheral positioning arises during development (Roman and Gomes, 2018) and is primarily due to the centre region of the muscle fibre being “crowded” with contractile material, leaving little space for myonuclei (Roman and Gomes, 2018, Roman et al., 2017). The peripheral placement may also facilitate their role as mechanosensors in muscle fibres (Van Ingen and Kirby, 2021), be related to vascular organization (Sequeira et al., 2024, Ralston et al., 2006), and contribute to proper regulation of the “myonuclear domain” (Bagley et al., 2023, Hansson et al., 2020). A preponderance of displaced and centralized myonuclei characterize degenerative and myopathic disease states such as muscular dystrophy (Grounds et al., 2014, Korb et al., 2024). Displaced myonuclei also manifest during conditions of extreme muscle stress such as chemical injury and severe trauma (Buckley et al., 2022, Pizza and Buckley, 2023, Hastings et al., 2020). Since muscle stem cells – or satellite cells – are indispensable for muscle regeneration (McCarthy et al., 2011, Lepper et al., 2011, Murphy et al., 2011, Sambasivan et al., 2011), the presence of displaced myonuclei is classically attributed to the activities of these cells (Pizza and Buckley, 2023). Regenerating muscle fibres are initiated by satellite cell fusion resulting in displaced or centralized myonuclei in myotubes that may ultimately relocate peripherally when the muscle fibre matures (Pizza and Buckley, 2023, Korb et al., 2024). Satellite cells are widely accepted as the explanation for displaced and centralized myonuclei in nearly all contexts, but there is a growing understanding that resident myonuclei are more mobile than previously appreciated in mature muscle fibres (Bagley et al., 2023, Roman and Muñoz-Cánoves, 2022). Recent evidence shows that resident non-satellite cell-derived myonuclei migrate to the site of injury after focal membrane damage to support repair in mature muscle fibres (Roman et al., 2021). Furthermore, displaced myonuclei are a feature of muscle fibres after minimally injurious voluntary exercise (Dungan et al., 2019), appear concomitant with muscle aging independent from severe injury (Cristea et al., 2010, Roth et al., 2000), emerge during muscle loading in the absence of overt degeneration/regeneration (Murach et al., 2017), and are characteristic of denervation as well as certain non-degenerative myopathies (Grounds et al., 2014). Displaced myonuclei are lower in prevalence during mechanical loading in the absence of satellite cells relative to satellite cell replete muscle but not completely eliminated (McCarthy et al., 2011). These observations suggest that displaced myonuclei can originate from non-satellite cell sources. Against this background, we ask: what is the source of displaced myonuclei in healthy adult skeletal muscle?

To address this fundamental question, we used our recombination-independent muscle fibre-specific doxycycline-inducible murine genetic model of fluorescent myonuclear labelling, called HSA-GFP (<u>h</u>uman <u>s</u>keletal <u>a</u>ctin reverse tetracycline transactivator tetracycline response element histone 2b <u>g</u>reen fluorescent <u>p</u>rotein) (Iwata et al., 2018, Wen et al., 2021, von Walden et al., 2020, Murach et al., 2021, Murach et al., 2022). With this model, we can fluorescently-label myonuclei in the presence of doxycycline and the GFP signal will remain once the GFP transgene is inactivated following the removal of doxycycline. To encourage the appearance of displaced myonuclei in the absence of catastrophic damage, we utilized the synergist ablation mechanical overload (MOV) approach. MOV is a loading and lengthening stimulus for skeletal muscle that can cause the appearance of displaced myonuclei above levels observed in unperturbed muscle (Murach et al., 2017, McCarthy et al., 2011). Myonuclei of adult mice (∼10-month-old) were labelled with GFP prior to a washout period, then muscle was subjected to MOV for 3 or 7 days; sham operated mice served as controls (Figure 1A). We quantified displaced myonuclei (defined as being detached from the sarcolemma on the interior of the muscle fibre) that were derived from pre-existing resident myonuclei (GFP+) versus those that were acquired from an outside source such as satellite cell fusion (GFP-) during MOV. We delivered 5-Ethynyl-2’-deoxyuridine (EdU) during MOV to quantify DNA synthesis and infer stem cell fate and analysed displaced myonuclei according to myosin heavy chain (MyHC) fibre type and overall fibre size.

**Figure 1.**
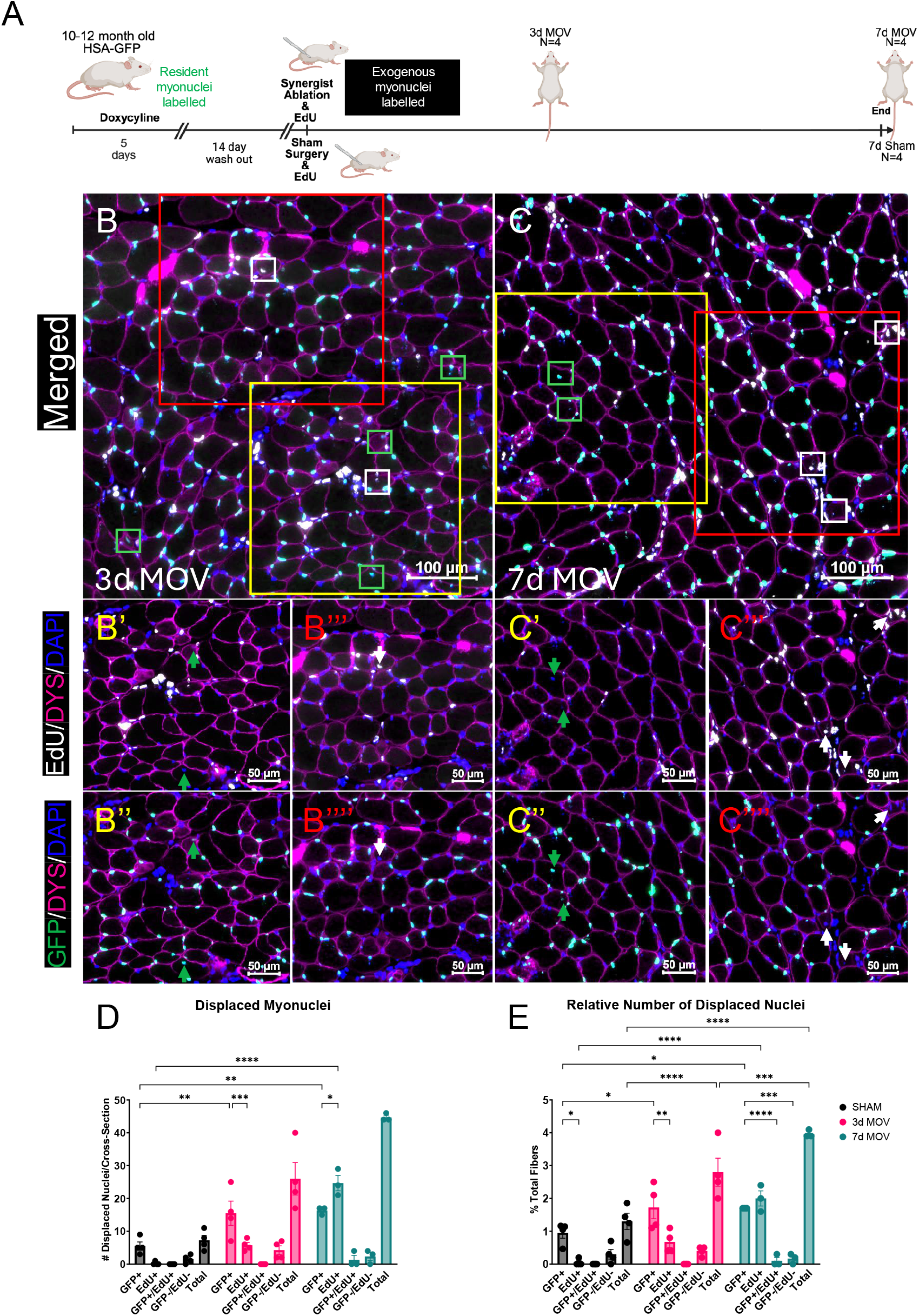
(A) Study design where 10-month-old HSA-GFP mice (M/F=6/6) were treated with doxycycline to label myonuclei for 5 days followed by a 14-day wash-out. Synergist ablation mechanical overload (MOV) surgery to overload the plantaris muscle was performed and mice were treated with 5-ethynyl-2’-deoxyuridine (EdU) in drinking water to assess DNA synthesis during overload. MOV muscles were collected after 3 days (N=4; M/F=2/2) and 7 days (N=4; M/F=2/2) with sham controls collected at 7 days (N=4; M/F=2/2, images not shown). 3-day MOV (B-B’’’’), and 7-day MOV (C-C’’’’) were probed for dystrophin (pink) and DAPI (blue) with GFP resident myonuclei in green and exogenous myonuclei (EdU+/GFP-) in white. Quantification of the absolute and relative (percentage) of total fibres with GFP+ displaced myonuclei (discernibly within but not adjacent to the dystrophin boarder) is reported in D&E. Data are presented as Mean ± SEM. (*p≤0.05; \*\**p*≤0.01, \*\*\**p*≤0.001,\*\*\*\**p*≤0.0001; 2-way ANOVA Treatment x Nuclei Label). EdU – 5-ethynyl-2’-deoxyuridine, DYS – dystrophin, GFP – green fluorescent protein. White boxes and arrows = EdU+, green boxes and arrows = GFP+

## Results and Discussion

### Displaced myonuclei have differential contributions from resident versus exogenous sources depending on MOV duration

Representative images of GFP+ (EdU-) and EdU+ (GFP-) displaced myonuclei (all DAPI+ and within the dystrophin border) are shown in Figures 1B&C (3d & 7d MOV), 1B-B’’’ (3-day MOV), and 1C-C’’’ (7-day MOV). The sham condition was time-matched to the 7-day MOV time point. Displaced myonuclei in the sham condition were very infrequent but almost exclusively GFP+ (images not shown); this means they were resident myonuclei that migrated during the 7 days after sham surgery or were already in a non-peripheral location during the doxycycline labelling period (Figure 1D&E). We found a significantly higher proportion of fibres with displaced myonuclei at 3 days of MOV relative to sham (∼25 fibres per cross section, or ∼3% of all fibres, *p*<0.0001) (Figure 1D&E). Of those displaced myonuclei after 3 days of MOV, a larger proportion were GFP+ versus EdU+ (Figure 1D&E). Resident myonuclear migration is therefore an early-emerging feature of the muscle adaptive response to MOV. By 7 days of MOV, the contribution of EdU+ myonuclei (exogenous, presumably from satellite cells) was similar to that of GFP+ displaced myonuclei (∼20 fibres per cross section of each type, ∼4% of all fibres, *p*=0.009) (Figure 1D&E). Recent studies show that satellite cells can fuse to muscle fibres during the early phase of MOV without a prior round of cell division (Ismaeel et al., 2024, Goh et al., 2025). We therefore quantified GFP-/EdU-displaced myonuclei, which could be indicative of direct satellite cell fusion. There were slightly more double-negative events after 3 days of MOV versus sham (Figure 1D&E). The overall number of displaced GFP+/EdU+ was negligible across conditions, suggesting that displaced resident myonuclei did not undergo measurable DNA synthesis over a 3-to-7-day MOV duration according to our detection methods. It is worth mentioning that the majority of displaced myonuclei that we observed were not in a centralized location in the muscle fibre. Perhaps more displaced myonuclei would ultimately adopt a central position with longer MOV duration. Centralized myonuclei are classically associated with a satellite cell-mediated regenerative response but could also potentially be related to cell-autonomous muscle healing processes, as described by Roman et al. (Roman et al., 2021). Myonuclear positioning near the muscle fibre membrane, but not abutting it, as we mostly observe here may be a feature of horizontal myonuclear movement along the sarcolemma versus vertical movement toward the centre of the fibre. Additional time-resolved experimentation is required to clarify the significance of varied positioning of displaced myonuclei.

### Fiber type-specific differences in displaced myonuclei during MOV

Since the plantaris muscle has a mix of MyHC IIb and non-IIb muscle fibres, we determined the occurrence of displaced myonuclei according to adult myosin fibre type. Representative images of fibre type-specific displaced myonuclei after MOV are shown in Figure 2A&A’ (3d) and 2B&B’ (7d). After 3 days of MOV, there was a similar proportion of MyHC IIb+ and IIb-muscle fibres with displaced GFP+ myonuclei (Figures 2C&D). GFP-displaced myonuclei were lower in proportion relative to GFP+ after 3 days of MOV, but the contribution from both sources was similar in MyHC IIb+ and IIb-fibres (Figures 2C&D). At 7 days of MOV, MyHC IIb+ fibres had more displaced myonuclei than MyHC IIb-fibres. These data generally align with observations of fibre type-specific sensitivity to loading-induced muscle damage in a given muscle (Macaluso et al., 2012, Vijayan et al., 2001), which is perhaps due to ultrastructural differences between fibre types (Qaisar et al., 2016). The relative contribution of GFP+ versus GFP-displaced myonuclei after 7 days of MOV was similar across fibre types (Figure 2C&D).

**Figure 2.**
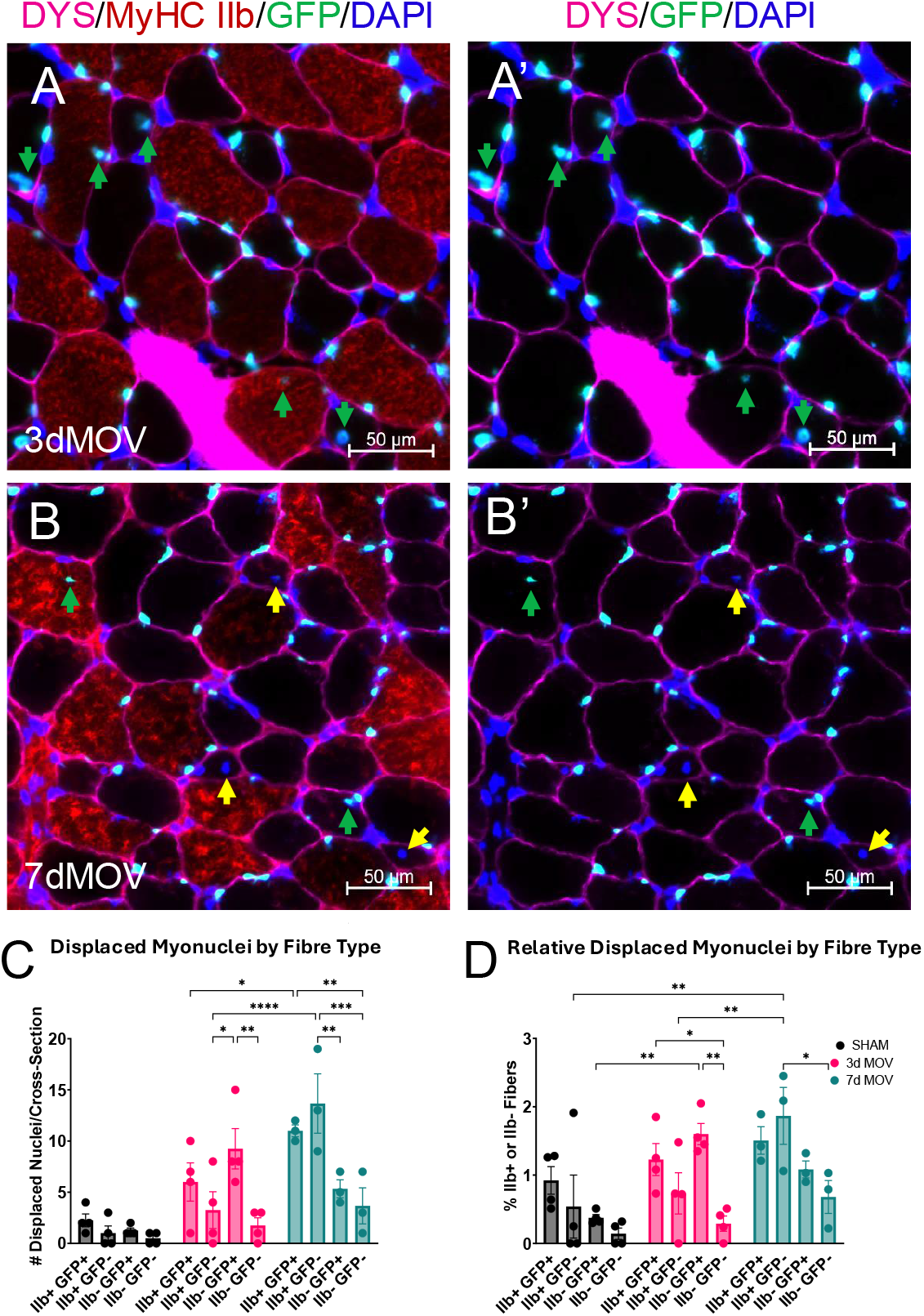
Immunohistochemical representative images of the plantaris muscle for fibre type-specific quantification of displaced myonuclei for 3-day MOV (A&A’) and 7-day MOV (B&B’). Images show dystrophin (pink), myosin heavy chain IIb (MyHC IIb, red), GFP resident myonuclei (green), and DAPI nuclei (blue). Quantification of displaced myonuclei according to fibre type and condition is reported in C&D. Data are presented as Mean ± SEM and is reported relative to each respective fibre type. (\**p*≤0.05; \*\**p*≤0.01, \*\*\**p*≤0.001,\*\*\*\**p*≤0.0001; 2-way ANOVA Treatment x Nuclei Label). MOV – mechanical overload, EdU – 5-ethynyl-2’-deoxyuridine, DYS – dystrophin, GFP – green fluorescent protein. Yellow arrows – GFP-, Green arrows – GFP+

### The origin of displaced myonuclei during MOV is fibre size-dependent

We next asked whether displaced GFP+ versus GFP-myonuclei varied according to muscle fibre size. In sham mice, relatively infrequent GFP+ displaced myonuclei tended to be dispersed across fibre sizes (Figure 3A&B). At 3 days of MOV, a pattern began to emerge where larger muscle fibres featured displaced GFP+ myonuclei and smaller fibres tended to have GFP-myonuclei (Figure 3C&D). After 7 days of MOV, it became clear that the largest fibres mostly contained displaced GFP+ myonuclei, and smaller fibres (<1,000 µm^2^) tended to have displaced GFP-myonuclei (Figure 3E&F). At the 3-day time point, our observations may be driven by displaced EdU+/GFP-myonuclei emerging in inherently smaller MyHC IIA fibres (see Figure 2). The early stage of muscle regeneration would result in the appearance of smaller calibre fibres with centralized myonuclei, but the 3-day time point is likely too early for muscle fibre regeneration to have progressed appreciably. By 7 days, however, the appearance of smaller muscle fibres with GFP-displaced myonuclei could be the result of nascent muscle fibre formation from a satellite cell-mediated regenerative response. To this point, there was one 7-day MOV sample with the appearance of widespread mononuclear cell infiltration and areas lacking muscle fibres with GFP+ myonuclei (i.e. muscle fibre degeneration), and a high density of EdU+ mononuclear cells in interstitial spaces. Within the infiltrated regions, very small muscle fibres tended to be positive for embryonic myosin heavy chain (eMyHC, a sign of muscle fibre regeneration) and have GFP-/EdU+ displaced myonuclei (Figure 4A-A’’’). In the other 7-day MOV muscles (n=3), less frequent and more regionalized eMyHC+ muscle fibres either contained no displaced myonuclei, or the displaced myonuclei in these fibres were majority EdU+ (Figure 4B-D). Overall, our data suggest that larger muscle fibres tend to experience resident myonuclear migration (GFP+) that results in displaced myonuclei, whereas smaller and/or regenerating muscle fibres have displaced myonuclei due to stem cell contributions (GFP-or EdU+).

**Figure 3.**
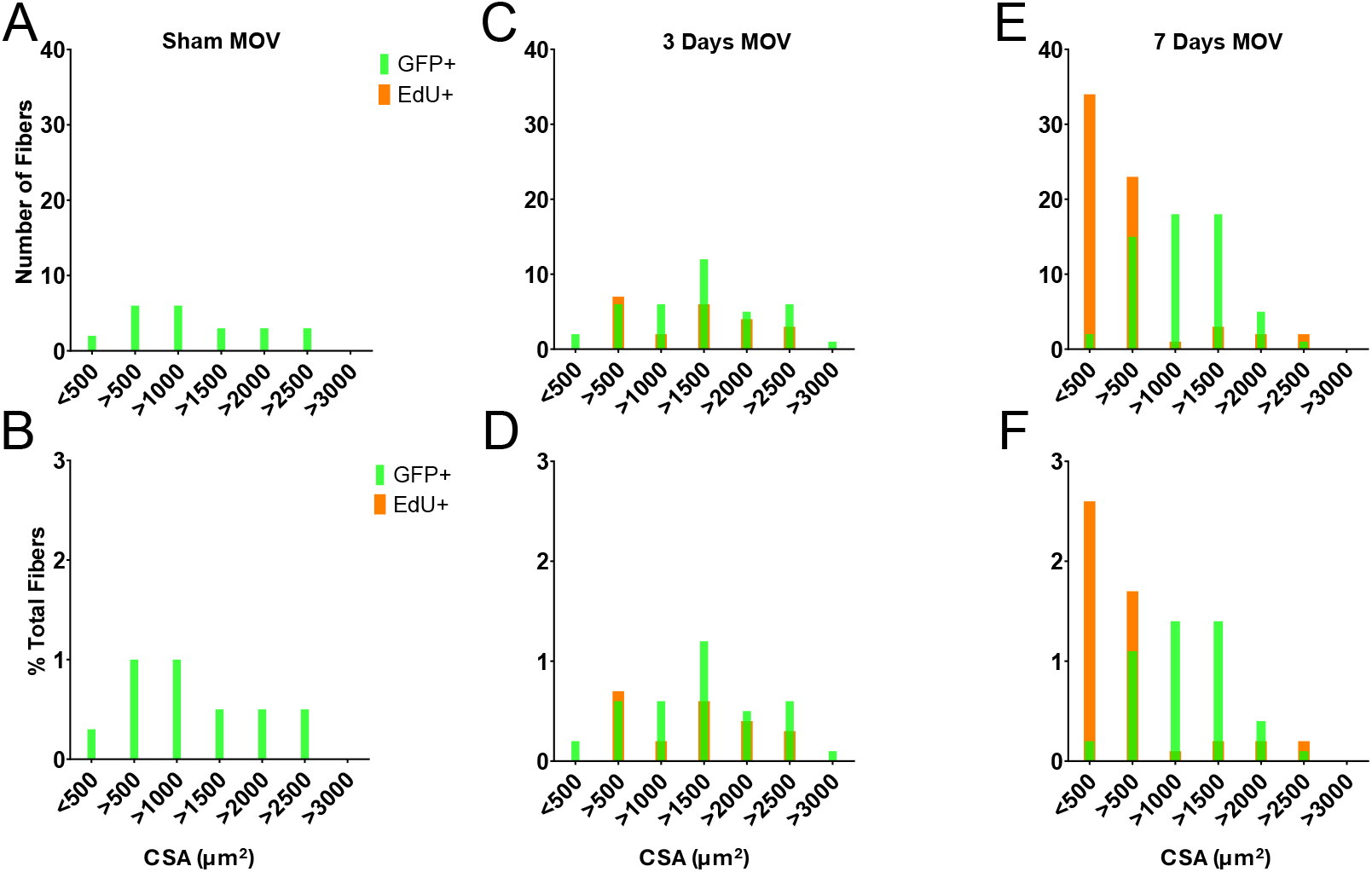
Frequency distributions show the occurrences of GFP+ (green) and EdU+ (white) displaced nuclei according to fibre cross sectional area (A,C,E). Panels B, D, & F show these frequencies as a proportion of total fibre counts on entire plantaris cross sections. Data are presented as Mean ± SEM and frequency distributions. (\**p*≤0.05; 2-way ANOVA Treatment x Nuclei Label). MOV – mechanical overload, EdU – 5-ethynyl-2’-deoxyuridine, CSA – cross sectional area, DYS – dystrophin, GFP – green fluorescent protein, MyHC IIb – myosin heavy chain type IIb

**Figure 4.**
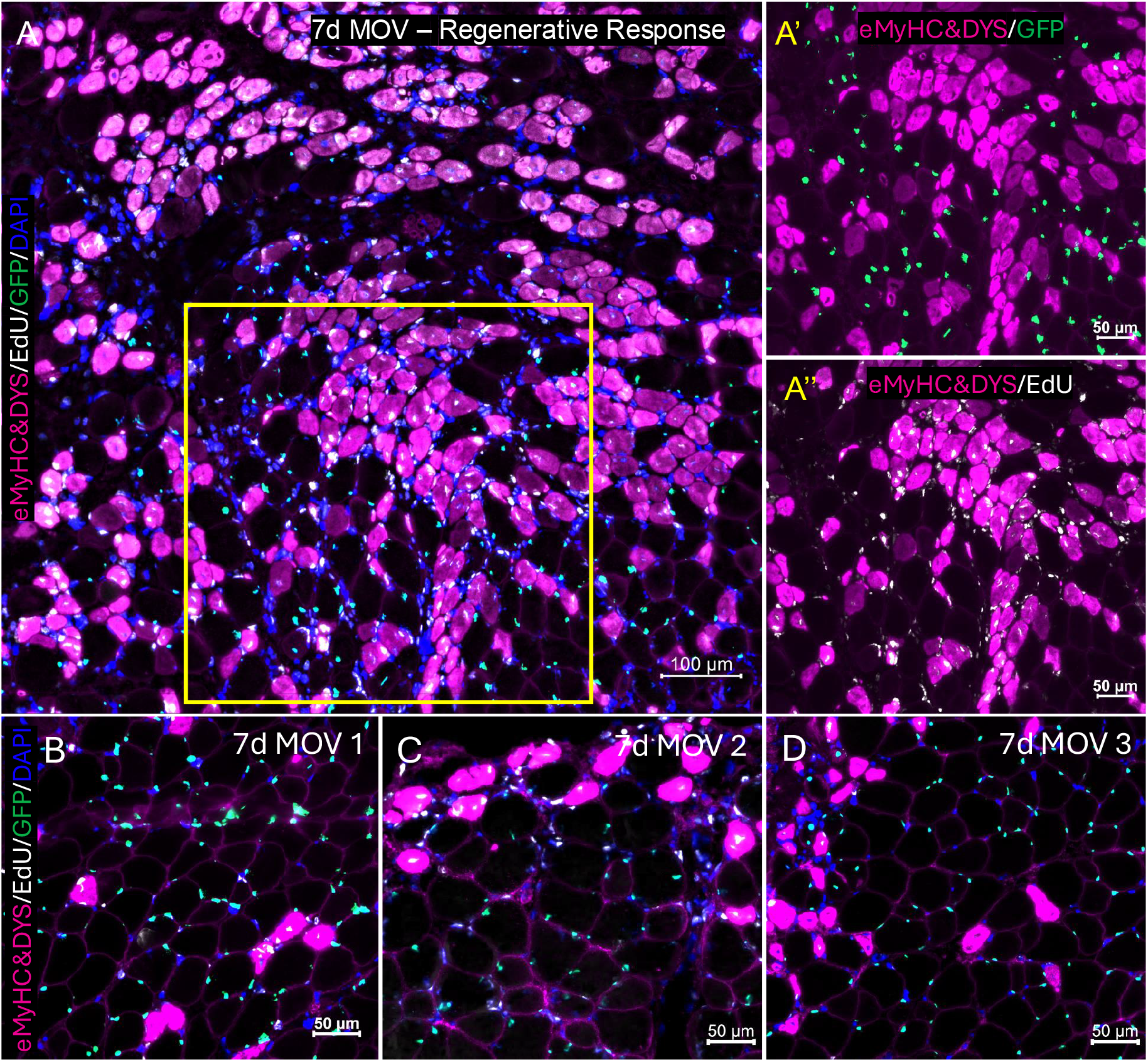
Image of a case study of degeneration/regeneration during MOV showing dystrophin (pink), EdU labelled nuclei (white), and DAPI (blue) with GFP resident myonuclei in green (A-A’’). Several fibres in the region of interest (yellow box in A) show displaced GFP+ myonuclei in both eMyHC+ and eMyHC-fibres surrounded by smaller fibres where displaced myonuclei are GFP-. Panels B-D show EdU+ displaced myonuclei occurring in the smaller eMyHC+ fibres of the normal 7d MOV muscles, which emerged regionally on the muscle cross-section. eMyHC - embryonic myosin heavy chain

### Study limitations

There are limitations to our investigation that should be considered. While GFP labelling in the doxycycline-inducible HSA-GFP model is highly specific to myonuclei, there is very rare off-target labelling of satellite cells (Iwata et al., 2018). It is possible that GFP+/EdU-displaced myonuclei are the result of errantly labelled GFP+ satellite cells, and that these same cells were the ones that fuse without a prior division (thus lacking EdU) (Ismaeel et al., 2024, Goh et al., 2025). However, the overall abundance of GFP+/EdU-displaced myonuclei in context with the similar abundance of GFP-/EdU+ displaced myonuclei makes a potentially small amount off-target satellite cell labelling an unlikely explanation for the appearance of all displaced GFP+ myonuclei. We delivered EdU in drinking water to identify DNA synthesis and cell fate. The half-life of EdU in rodents *in vivo* is short (a matter of hours) (Solius et al., 2021). A more rigorous approach that would label more EdU+ events is to use a mini osmotic pump that delivers EdU continuously throughout MOV (Ismaeel et al., 2024, Goh et al., 2025). Recent evidence suggests that resident myonuclei can synthesize DNA, and that myonuclear DNA synthesis is elevated during MOV (Borowik et al., 2022, Borowik et al., 2024). The usage of a mini osmotic pump approach to deliver EdU could potentially capture DNA synthesis in displaced resident myonuclei assuming off-target GFP labelling of satellite cells or alternative progenitors (Liu et al., 2017, Flynn et al., 2023, Dellavalle et al., 2011, Mitchell et al., 2010)-followed by proliferation and fusion-does not explain the result. Notwithstanding, a preponderance of EdU+ events in regenerating areas of muscle during MOV in our experiments (see Figure 4A-A’’) confirms that our labelling was robust. The proportion of muscle fibres with displaced myonuclei is relatively low in the MOV condition (up to 4% of all fibres), but it is important to consider that our analyses were performed on 7 µm sections. If extrapolated to the entire length of the plantaris muscle, this would equate to an appreciable number of displaced myonuclei.

### Perspectives

We previously associated the appearance of a displaced myonucleus - in the absence of an overt regenerative response - to the occurrence of what appeared to be fibre “splitting” during MOV (Murach et al., 2017). A provocative hypothesis is that displaced resident (non-stem cell-derived) myonuclei could be a biomarker for, or even a contributor to the rare “splitting” of an adult muscle fibre that might occur during MOV (Murach et al., 2019). Future investigations may explore this possibility. The results from the current study build on our preliminary observations of displaced resident myonuclei with muscle damage (Murach et al., 2020) and provide new perspectives on the source, timing, myosin type, and size characteristics of fibres containing non-peripheral myonuclei in adult muscle fibres. Given the role resident myonuclear migration plays in muscle fibre membrane repair (Roman and Muñoz-Cánoves, 2022, Roman et al., 2021, Bagley et al., 2023), our work provides motivation to study the origin and function of displaced myonuclei across different conditions such as muscle disease and aging and re-evaluate our understanding of regenerating muscle fibres based solely on centralized myonuclear placement (Grounds et al., 2014). The molecular characteristics of resident versus exogenous displaced myonuclei should also be explored. More granular information on the source and function of displaced myonuclei in adult skeletal muscle could further illuminate etiology and treatment apporaches for muscle damage and pathology, as well as myonuclear contributions to exercise adaptation (Koopmans et al., 2023).

## Methods

### Ethical approval

All animal procedures were approved by the IACUC of the University of Arkansas, Fayetteville. The human skeletal actin reverse tetracycline transactivator tetracycline response element histone 2b green fluorescent protein (HSA-GFP) mice were generated by our laboratory. Mouse pups were genotyped as previously described (Iwata et al., 2018). Mice were housed in a temperature and humidity-controlled facility with 12:12h light-dark cycle with food and water provided ad libitum. Animals were sacrificed by being placed under general anaesthesia with isoflurane followed by cervical dislocation.

### Experimental Design

The experimental design is illustrated in Figure 1A. 10 month old mice (N=12; M/F=6/6) were assigned to either a 3-(n=4; M/F=2/2) or 7-day MOV (n=4; M/F=2/2) protocol with a 7-day time matched sham group (Sham) (n=4; M/F=2/2). One 7-day MOV mouse had the appearance of overt degeneration/regeneration and was treated as a “case study”, separate from the rest of the group (see Figure 4) and not included in other analyses. Mice were treated with 0.5mg/ml doxycycline and 2% sucrose for 5 days in drinking water to fluorescently label resident myonuclei. After a 14-day washout period, mice underwent our synergist ablation mechanical overload surgery where the lower 2/3 of the gastrocnemius and soleus complex were removed. Mice returned to normal ambulation within ∼24 hours. At the time of MOV surgery, mice were delivered an intraperitoneal “priming dose” of EdU (∼2 mg in water), then given 0.5 mg/ml EdU (Biosynth, NE08701) with 2% sucrose in drinking water that was refreshed every other day, similar to our prior approach (Murach et al., 2020).

### Histology/Immunohistochemistry

Immunohistochemistry (IHC) was carried out on whole plantaris muscles after embedding in optimal cutting temperature (OCT) compound and snap freezing in liquid nitrogen chilled isopentane as previously described by our us using appropriate antibodies (Murach et al., 2020). For assessment of displaced nuclei, plantaris muscles were cryosectioned at 7 µm on a Epredia Cryostar NX50 Cryostat and fixed with 4% paraformaldehyde and permeabilized in 0.5% Triton-X. EdU detection was carried out with CLICK-iT detection buffer for 30 minutes that contained 100mM Tris, 5mM Copper Sulphate, 100mM ascorbic acid, and TAMRA-Azide (Vector, CCT-AZ109-1). Anti-dystrophin primary was then applied and an appropriate secondary antibody followed by DAPI prior to mounting and imaging. For fibre type assessments, sections were incubated in primaries for MyHC IIb (BF-F3, DSHB) and dystrophin then the appropriate secondary antibodies (isotype-specific for MyHC IIb) before applying DAPI. Displaced myonuclei were defined as myonuclei that are not adjacent to the sarcolemma (dystrophin) and manually assessed in Zen software.

### Embryonic Myosin Immunohistochemistry

Separate sections were blocked with 10% normal horse serum and 1% BSA in PBS for 30 minutes at room temperature before applying eMyHC (F1.652, DSHB) and dystrophin primary antibody incubated at 4°C. Secondary antibody was then applied before post-fixing in 4% PFA and continuing with EdU detection as described in the previous section.

### Image Capture

All images were captured using an upright fluorescent microscope at 20X magnification (Zeiss Axiolmager M2, Oberkochen, Germany) where whole muscle sections were imaged using the mosaic function in Zeiss Zen 3.8.3 for Microsoft. Muscle fibre counts on whole muscle sections were assessed in MyoVision semi-automated analysis software. Displaced nuclei were manually counted in Zen software where fibres with displaced nuclei were then manually assessed for cross-sectional area (CSA) using the fibre tracing tool.

### Statistics

Displaced nuclei counts were analysed using GraphPad statistical software (Prism, version 10.6.0 for Windows). A 2-way ANOVA was performed on the GFP+, EdU+, GFP+/EdU+, and GFP-/EdU-displaced myonuclear counts per section and the percentage of nuclei relative to total fibre counts with a Tukey’s post hoc correction. A separate 2-way ANOVA on the displaced myonuclei per fibre type (type IIb vs non-IIb) with a tukey’s post hoc correction and significance set at p≤0.05. Finally, frequency distributions were collected and incrementally binned using the manually traced fibre CSA data from the Zeiss Zen software and personalized Excel macros. All figures were created using GraphPad.

## Acknowledgments

Thank you to Cory Dungan, PhD, of Baylor University for helpful discussions regarding EdU detection.

